# No circadian clock-gene circuit in the generation of cellular bioluminescence rhythm of *CaMV35S::PtRLUC* in duckweed

**DOI:** 10.1101/2022.05.12.491730

**Authors:** Emiri Watanabe, Tomoaki Muranaka, Shunji Nakamura, Minako Isoda, Shogo Ito, Tokitaka Oyama

## Abstract

Physiological circadian rhythms are coordinated in the plant body in an orderly manner. The coordination of time information has been studied from the aspects of cell–cell local coupling and long-distance communication between tissues. These studies were based on the idea that the behavior of the clock gene circuit represents the physiological rhythms. Here we report the cellular circadian rhythm of a bioluminescence reporter which is not governed by the clock gene circuit in the expressing cells. Using a dual-color bioluminescence monitoring system in *Lemna minor* transfected with the *AtCCA1::LUC*+ and *CaMV35S::PtRLUC* reporters, cellular bioluminescence rhythms with different free-running periods (FRPs) were detected in the same cells. Co-transfection experiments with the two reporters and a clock gene overexpressing effector revealed that the circadian properties of the *AtCCA1::LUC*+ rhythm, but not those of the *CaMV35S::PtRLUC* rhythm, were altered in the cells with a dysfunctional clock gene circuit. This indicates that the *AtCCA1::LUC*+ rhythm is a direct output of the cellular circadian clock while the *CaMV35S::PtRLUC* rhythm is not. After plasmolysis, the *CaMV35S::PtRLUC* rhythm disappeared while the *AtCCA1::LUC*+ rhythm persisted. The plant circadian system consists of both cell-autonomous rhythms and non-cell-autonomous rhythms that are unaffected by the cellular clock.

## Introduction

Many physiological phenomena in plants show circadian rhythms (Sweeney, 1987). The circadian clock is a cell-autonomous system composed of a number of clock genes that form transcription–translation feedback loops (TTFLs) (Nohales and Kay, 2016; Sanchez et al., 2020). In *Arabidopsis*, dawn-expressed *CIRCADIAN CLOCK-ASSOCIATED 1* (*CCA1*), *LATE ELONGATED HYPOCOTYL* (*LHY*), morning-to-evening-expressed *PSEUDO-RESPONSE REGULATOR* (*PRR*) family, evening-tonight-expressed *EARLY FLOWERING 3* (*ELF3*), *ELF4*, and *LUX ARRHYTHMO* (*LUX*) repress genes expressed during earlier phases. The F-box protein ZEITLUPE (ZTL) affects the TTFLs by temporal promotion of proteasomal degradation of PRR members. Disruption and/or overexpression of these clock genes impairs the circadian rhythm (Nohales and Kay, 2016; Sanchez et al., 2020). The TTFL oscillator governs the circadian rhythms of various physiological processes in plants, and mutations in clock genes alter these circadian rhythms (Somers et al., 1998; Xu et al., 2007).

Physiological rhythms with different free-running periods (FRPs) have been observed in plants (Nohales, 2021). For example, rhythms in cytosolic free calcium (Ca^2+^) levels have a different FRP from that of *Lhcb* gene expression in tobacco (Sai and Johnson, 1999). Differences in the FRPs of bioluminescence rhythms were observed in *Arabidopsis* seedlings expressing a firefly luciferase under circadian promoters *CHALCONE SYNTHASE* (*CHS*), *CHLOROPHYLL A/B BINDING PROTEIN* (*CAB*), *PHYTOCHROME B* (*PHYB*) and *CATALASE 3* (*CAT3*) (Hall et al., 2002; Michael et al., 2003; Thain et al., 2002). Tissue-specific modulation of the TTFL is considered to be a contributing factor to these different FRPs (McClung, 2019; Nohales, 2021). These distinct oscillators can bring about uncoupled rhythms between tissues/organs and influence the coordination of physiological processes at the organismal level.

In addition to tissue/organ-specific circadian rhythms, intracellular uncoupling of two circadian rhythms has also been reported in plant cells (Watanabe et al., 2021). Bioluminescence of a luciferase driven by a constitutive promoter showed a circadian rhythm in various duckweeds, and *CaMV35S::PtRLUC* (a modified click beetle red color luciferase gene) showed bioluminescence rhythms uncoupled from those of *AtCCA1::LUC*+ (a firefly yellow-green luciferase gene) in individual cells in *Lemna minor* (Muranaka et al., 2015; Watanabe et al., 2021). The uncoupled rhythms were detected using a dual-color bioluminescence monitoring system at a single-cell level, in which *AtCCA1::LUC*+ and *CaMV35S::PtRLUC* reporters were simultaneously transfected into the cells in the duckweed plant by particle bombardment. The intracellular uncoupling was observed as inconsistent phase relation and different FRPs under prolonged and constant light conditions. The *AtCCA1::LUC*+ bioluminescence rhythm was shown to be cell-autonomously generated under the control of the TTFL in individual cells, through functional analysis by co-transfecting an effector (overexpression, knockdown and knockout of clock genes) with the circadian reporter (Miwa et al., 2006; Okada et al., 2017; Serikawa et al., 2008). In duckweed plants, desynchronization of the cell-autonomous circadian clock occurs between cells in the same tissue under constant conditions, although weak cell-to-cell interaction influencing their phases is expected (Muranaka and Oyama, 2016; Ueno et al., 2022). However, it is unclear how the intracellular uncoupling between *AtCCA1::LUC*+ and *CaMV35S::PtRLUC* bioluminescence rhythms happens in individual cells, as the mechanism underlying the *CaMV35S::PtRLUC* bioluminescence rhythm is currently unknown.(Watanabe et al., 2021) Cell autonomy and the direct involvement of TTFL are questionable. In this study, we performed co-transfection using circadian reporters and clock gene overexpressing effectors in a dual-color bioluminescence monitoring system at a single-cell level. We also analyzed the effects of plasmolysis on the cellular circadian rhythms. We show that the *CaMV35S::PtRLUC* bioluminescence rhythm is non-cell-autonomously generated, without direct regulation by the clock-gene circuit.

## Results

### Differences in the free-running periods and the intercellular synchrony between *AtCCA1::LUC*+ and *CaMV35S::PtRLUC* bioluminescence rhythms

In order to directly compare circadian properties of the bioluminescence rhythms between the two reporters, we performed dual-color bioluminescence monitoring at a single-cell level (Watanabe et al., 2021). Circadian reporters with different luminescence colors, *AtCCA1::LUC*+ (yellow-green) and *CaMV35S::PtRLUC* (red), were introduced into *L. minor* cells (epidermal and mesophyll cells) through particle bombardment (Figure 1A). The bioluminescence of individual cells was observed using an EM-CCD camera with a filter exchanger containing green-pass and red-pass filters, and the cellular luminescence intensities of each reporter were reconstructed based on a set of filtered luminescence images (Figure 1B). A set of luminescence images was acquired every hour according to the time schedule shown in Figure 1C.

**Figure 1.**
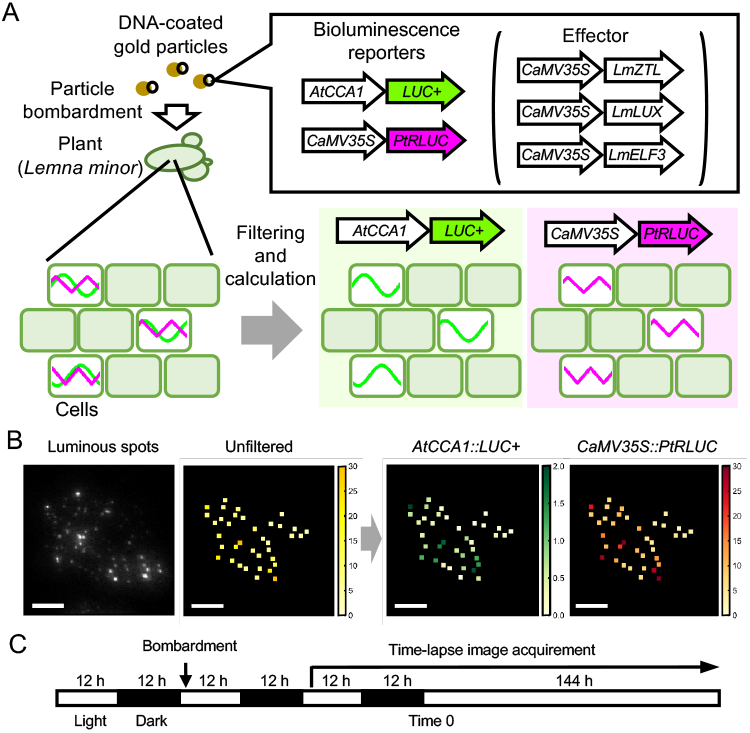
Outline of the dual-color bioluminescence monitoring experiments. (A) Schematic drawing of the measurement of *AtCCA1::LUC*+ and *CaMV35S::PtRLUC* bioluminescence at a single-cell level. Bioluminescence reporters were introduced into *Lemna minor* cells by particle bombardment. Filtered cellular bioluminescence was imaged and used for the calculation of luminescence intensities of each reporter. For overexpression experiments, an effector (*CaMV35S::LmZTL, CaMV35S::LmLUX* or *CaMV35S::LmELF3*) was co-transfected with the reporters. (B) An example of the reconstruction of cellular luminescence intensities [1000 photons/s] of each reporter. Luminescence intensities of individual cells are color coded. Scale bars are 1 mm. (C) Time schedule of the monitoring experiments. *L. minor* plants that had been cultured under 12 h light 12 h dark cycles were transfected by particle bombardment. The transfected plants were further cultured under 12 h light 12 h dark cycles, and then transferred to the imaging system. Open and closed boxes represent light and dark conditions, respectively. Time 0 is defined as the start of constant light during the timelapse imaging.

*AtCCA1::LUC*+ and *CaMV35S::PtRLUC* bioluminescence rhythms of 37 cells from the same frond under constant light are shown in Figure 2. As shown in Figure 2A and 2B, most cells showed circadian rhythms of both bioluminescence reporters. The FRP and relative amplitude error (RAE) of each luminescence trace were calculated using the fast Fourier transform non-linear least squares (FFT-NLLS) method (Plautz et al., 1997). The RAE value represents the degree of confidence of rhythmicity, ranging from 0 (complete sine-fitting) to 1 (arrhythmic). In this study, bioluminescence traces with RAE < 0.2 were defined as rhythmic, and the rest were defined as arrhythmic. Every cell showed *CaMV35S::PtRLUC* rhythm, while 33 cells (89%) showed *AtCCA1::LUC*+ rhythm (Table S1). Although the bioluminescence traces of *CaMV35S::PtRLUC* in all cells were judged to be rhythmic, the *CaMV35S::PtRLUC* rhythm looked lower in amplitude than the *AtCCA1::LUC*+ rhythm (Figure 2A and 2B). FRPs of *AtCCA1::LUC*+ and *CaMV35S::PtRLUC* rhythms were distributed differently, with values of 25.5 ± 0.8 h and 24.7 ± 0.6 h, respectively (Figure 2C and Table S1). The FRP of the *CaMV35S::PtRLUC* rhythm was significantly shorter than that of the *AtCCA1::LUC*+ rhythm in the same set of cells in a frond (Welch *t*-test, *p*=3.8×10^-13^). The shorter FRP of *CaMV35S::PtRLUC* rhythm was also observed in *L. minor* plants that had grown under constant light without experiencing an entrainment dark stimulus (Watanabe et al., 2021).

**Figure 2.**
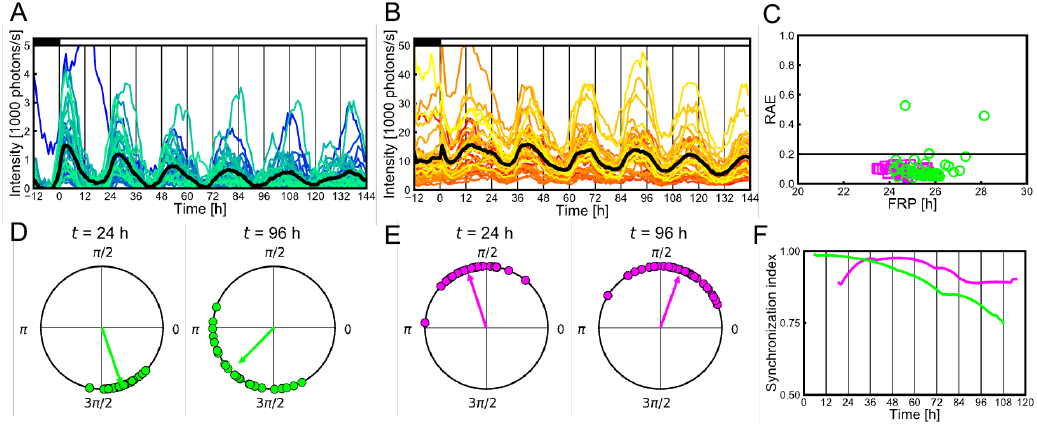
Cellular circadian properties of *AtCCA1::LUC*+ and *CaMV35S::PtRLUC* rhythms in the dual-reporter experiment. A total of 37 cells in a frond were analyzed. (A and B) Cellular *AtCCA1::LUC*+ (A) and *CaMV35S::PtRLUC* (B) bioluminescence rhythms. The black line represents mean luminescence intensities. Close and open boxes represent light and dark conditions, respectively. (C) The free-running periods (FRPs) and the relative amplitude errors (RAEs) of individual cellular rhythms. Green circles and magenta squares represent *AtCCA1::LUC*+ and *CaMV35S::PtRLUC*, rhythms, respectively. Cellular bioluminescence traces with RAE < 0.2 are defined as rhythmic. (D and E) Plots of the phases of *AtCCA1::LUC*+ rhythm (D) and *CaMV35S::PtRLUC* rhythm (E) on the unit circle at 24 h and 96 h, respectively. The peak right before 24 h or 96 h is referred to as phase 0 [rad]. The vector represents the vector mean of each plot on the unit circle. The vector length and the angle represent the synchronization index (SI) and a representative phase, respectively, at each time. (F) Temporal changes in the SIs of *AtCCA1::LUC*+ (green) and *CaMV35S::PtRLUC* (magenta) rhythms.

Figures 2D and 2E show the phase distributions and degrees of synchrony (synchronization index: SI) at t=24 h and 96 h for *AtCCA1::LUC*+ rhythms and *CaMV35S::PtRLUC* rhythms, respectively. The SI of *AtCCA1::LUC*+ rhythm was 0.96 at t=24 h and decreased to 0.66 at t=96 h. The SI of *CaMV35S::PtRLUC* at t=24 h was 0.85, which was almost the same as that at t=96 h (0.79). The SI of *AtCCA1::LUC*+ rhythm continuously decreased during the constant light condition, while that of *CaMV35S::PtRLUC* rhythm was stable (Figure 2F). Previous studies found that desynchronization of *AtCCA1::LUC*+ cellular rhythms occurred under constant light condition in duckweed plants that had been entrained.(Muranaka and Oyama, 2016; Ueno et al., 2022) The higher synchrony of *CaMV35S::PtRLUC* cellular rhythms in fronds with asynchronous *AtCCA1::LUC*+ cellular rhythms has also been reported in plants that had not experienced entrainment (Watanabe et al., 2021).

In *L. minor* transgenic plants carrying an *AtCCA1::LUC*+ reporter, the circadian behaviors of bioluminescence showed a variation among fronds.(Ueno et al., 2022) To investigate variations in cellular circadian behaviors of *AtCCA1::LUC*+ and *CaMV35S::PtRLUC* rhythms among fronds, we studied four fronds (including that shown in Figure 2) through dual-color bioluminescence monitoring (dual#1-#4; Table S1) and three fronds for each of the reporters through single reporter bioluminescence monitoring (sCCA1#1-#3, s35S#1-#3; Table S2). The mean FRPs of *AtCCA1::LUC*+ rhythm determined by dual-color monitoring were 25.3-27.4 h, and those of *CaMV35S::PtRLUC* rhythm were 24.0-25.0 h (Figure 3A). In three out of the four fronds, the FRPs of *CaMV35S::PtRLUC* rhythm were significantly shorter than those of *AtCCA1::LUC*+ rhythm. This was also confirmed by comparing FRPs of the two rhythms in individual cells; 97 out of 116 cells showed shorter FRPs in case of *CaMV35S::PtRLUC* rhythm (Figure 3B). Interestingly, the *CaMV35S::PtRLUC* rhythm (s^2^ = 0.67) showed lower FRP variation between cells than the *AtCCA1::LUC*+ rhythm (s^2^ = 2.60) (*F* test, *p* = 3.2×10^-12^). The FRP variation is linked to desynchronization of cellular rhythms in a frond.(Muranaka and Oyama, 2016) In fact, the SI of *AtCCA1::LUC*+ rhythm in each frond continuously decreased while that of *CaMV35S::PtRLUC* was stable (Figure 2F and S1A). These differences in cellular circadian properties between the *AtCCA1::LUC*+ and *CaMV35S::PtRLUC* rhythms were confirmed through single reporter bioluminescence monitoring (Table S2, Figure S1BC). With respect to the phase relation, the mean peak phases of the bioluminescence rhythms showed higher variation between fronds in dual-reporter experiments (*AtCCA1::LUC+*, 4.307-5.204 rad; *CaMV35S::PtRLUC*, 0.334-1.890 rad) than those in single reporter experiments (*AtCCA1::LUC+*, 5.284-5.654 rad; *CaMV35S::PtRLUC*, 0.706-1.087 rad) (Tables S1 and S2). The higher phase variation in the dual-reporter experiments might be due to noise amplification in the process of reconstruction for luminescence intensities of both reporters.

**Figure 3.**
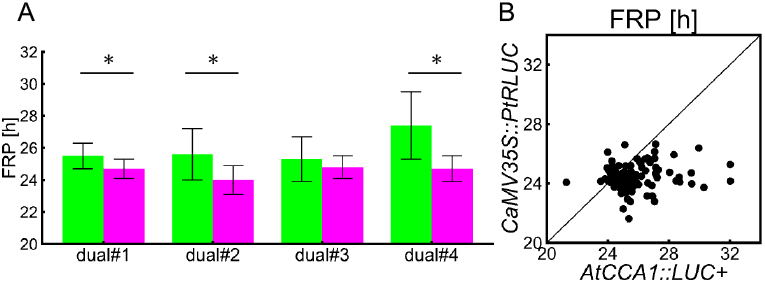
Difference in the free-running periods (FRPs) between *AtCCA1::LUC*+ and *CaMV35S::PtRLUC* bioluminescence rhythms. Cellular rhythms detected in the four experiments (dual#1: 33 cells, dual#2: 50 cells, dual#3: 18 cells, dual#4: 15 cells) were analyzed. (A) FRPs of *AtCCA1::LUC*+ or *CaMV35S::PtRLUC* rhythms in each experiment. Data are expressed as mean ± SD. Asterisk denotes statistically significant difference (Welch’s *t* test, *p* < 0.01). (B) A scatterplot of the FRPs between *AtCCA1::LUC*+ and *CaMV35S::PtRLUC* rhythms. FRPs of the 116 cells showing the circadian rhythms of both reporters in the four experiments are plotted. The dashed line represents where the FRPs are identical in both rhythms.

In summary, we noted the shorter period and higher synchrony of the *CaMV35S::PtRLUC* rhythm than the *AtCCA1::LUC*+ rhythm in the same frond through dual-color bioluminescence monitoring at a single cell level. The differences in the FRPs and the intercellular synchrony between the two bioluminescence rhythms indicate differences in the mechanisms of oscillation generation between them.

### Robustness of *CaMV35S::PtRLUC* bioluminescence rhythm against clock gene overexpression at the single cell level

We co-transfected the overexpressing effector of each of the three *L. minor* clock gene homologs, *LmZTL, LmLUX*, and *LmELF3*, with the dual reporters to examine their effects on the *AtCCA1::LUC*+ and *CaMV35S::PtRLUC* bioluminescence rhythms in the same cell (Table S1 and Figure S2). In *Arabidopsis*, it was reported that moderate and strong overexpression of *ZTL* caused shortening of FRP and loss of circadian rhythms, respectively (Somers et al., 2004). In *L. minor, LmZTL* overexpression caused a destabilization of *AtCCA1::LUC*+ rhythm (Table S1). When RAE<0.2 was used as a criterion for rhythmic cells, only 21% of the analyzed cells showed circadian rhythms in the experiment dual-ZTLox#1 (Figure 4A and 4C). Without the *CaMV35S::LmZTL* effector, most cells showed *AtCCA1::LUC*+ rhythm (Figure 2C and Table S1). This indicated that overexpression of *LmZTL* disrupted the clock gene circuit in the transfected cells. By contrast, overexpression of *LmZTL* did not affect the *CaMV35S::PtRLUC* rhythm in the analyzed cells. Every cell showed a waveform rhythm similar to that observed in experiments without the *CaMV35S::LmZTL* effector (Figures 2B and 4B). The mean FRP was 24.1 ± 0.6 h in the experiment dual-ZTLox#1, which was similar to that without the effector (Table S1). It is noteworthy that *AtCCA1::LUC*+ bioluminescence traces in rhythmic cells (RAE<0.2) in case of dual-ZTLox#1 showed a waveform and phase similar to those of the *CaMV35S::PtRLUC* bioluminescence rhythm (Figures 4A and 4B). This *CaMV35S::PtRLUC*-type bioluminescence rhythm exhibited by *AtCCA1::LUC*+ was also observed in the single reporter experiment with *LmZTL* overexpression (Figure S3). This indicates that the *AtCCA1::LUC*+ transfected cells are capable of producing *CaMV35S-type* bioluminescence rhythm when the clock-gene circuit is disrupted by the overexpression of *LmZTL*. Those cells in which the TTFL oscillator is disrupted may therefore have the potential to produce the *CaMV35S::PtRLUC*-type bioluminescence rhythm with any luciferase reporter in the co-transfection experiments.

**Figure 4.**
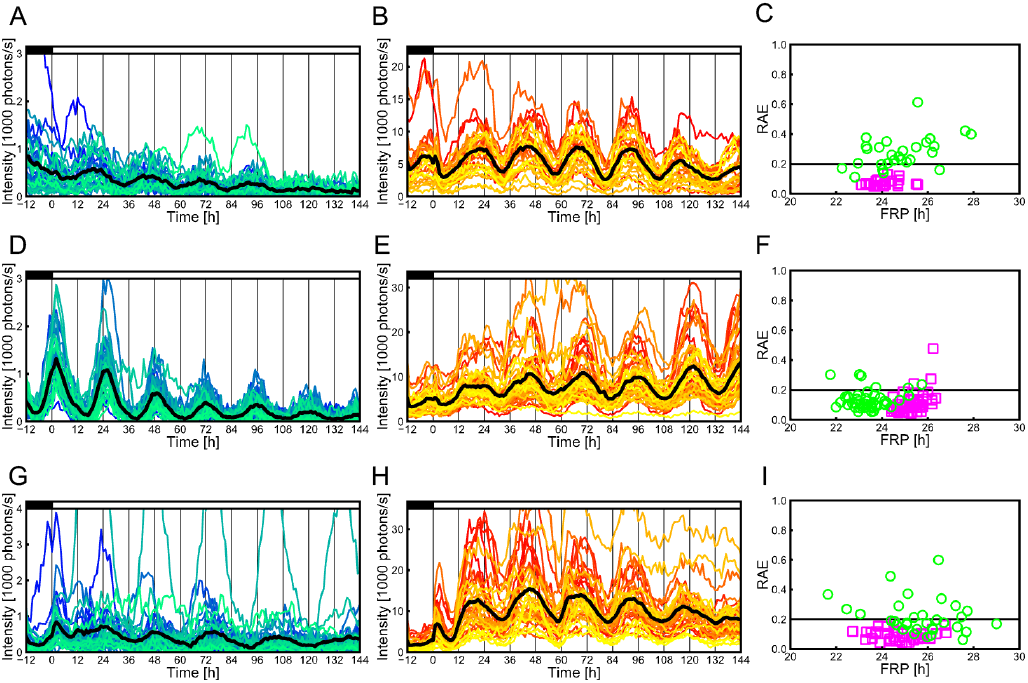
Cellular circadian properties of *AtCCA1::LUC*+ and *CaMV35S::PtRLUC* bioluminescence traces in clock-gene overexpression experiments. *CaMV35S::LmZTL* (dual-ZTLox#1, 29 cells) (A, B, C), *CaMV35S::LmLUX* (dual-LUXox#1, 48 cells) (D, E, F) and *CaMV35S::LmELF3* (dual-ELF3ox#1, 38 cells) (G, H, I) were co-transfected with the dual-color reporters. Cellular *AtCCA1::LUC*+ (A, D, G) and *CaMV35S::PtRLUC* (B, E, H) bioluminescence traces are lined in color and a black line in each panel represents mean luminescence intensities. Close and open boxes represent light and dark conditions, respectively. (C, F, I) The free-running periods (FRPs) and the relative amplitude errors (RAEs) of individual cellular rhythms. Green circles and magenta squares represent *AtCCA1::LUC*+ and *CaMV35S::PtRLUC* bioluminescence traces, respectively. Cellular bioluminescence traces with RAE < 0.2 are defined as rhythmic.

In transgenic *Arabidopsis* overexpressing *LUX* (*PCL*), dampening of the rhythmic expression of clock-genes has been reported.(Onai and Ishiura, 2005) In *L. minor, LmLUX* overexpression shortened the FRP of *AtCCA1::LUC*+ rhythm with an amplitude similar to that in the experiment without the effector (Figures 4D, 4F and Table S1). By contrast, *LmLUX* overexpression did not affect the *CaMV35S::PtRLUC* rhythm in the analyzed cells. Every cell showed a waveform and FRP similar to those in experiments without the effector (Figures 2D, 4E, 4F and Table S1). Consequently, *LmLUX* overexpression shortened the FRPs of the *AtCCA1::LUC*+ rhythm compared with those of the *CaMV35S::PtRLUC* rhythm in most of the analyzed cells (Figures S2 and S4). This indicated that *LmLUX* overexpression disrupted the clock gene circuit in the transfected cells without affecting the *CaMV35S::PtRLUC* rhythm.

Overexpression of *LgELF3* in the cells of *Lemna gibba* resulted in perturbation of light entrainment, and an increase in the number of cells with unstable rhythms and longer FRPs (Okada et al., 2017). In *L. minor*, overexpression of *LmELF3* caused unstable *AtCCA1::LUC*+ cellular rhythms and longer FRPs in dual-reporter experiments (Figure 4G, 4I and Table S1). By contrast, cellular *CaMV35S::PtRLUC* bioluminescence showed a stable rhythm and no alteration of FRP (Figure 4H, 4I and Table S1). This indicated that overexpression of *LmELF3* modified the clock gene circuit in the transfected cells without affecting the *CaMV35S::PtRLUC* rhythm. In single-reporter experiments, many cells co-transfected with *LmELF3* overexpressing effector showed unstable *AtCCA1::LUC*+ rhythm but stable *CaMV35S::PtRLUC* rhythm (Table S2). These results suggested that *LmELF3* overexpression impaired the clock gene circuit in the transfected cells that maintained the *CaMV35S::PtRLUC* rhythm. Results of the overexpression of the three clock genes, therefore, indicate that the *CaMV35S::PtRLUC* bioluminescence rhythm is generated on a cellular level without direct regulation by the clock gene circuit in the same cell.

### Disruption of the *CaMV35S::PtRLUC* bioluminescence rhythm in a hypertonic medium

While the co-transfection of a clock-gene overexpressing effector affects the circadian clock in the luciferase-expressing cells, most cells not transfected by particle bombardment are unlikely to be affected by the effector. Therefore, it raises the possibility that the *CaMV35S::PtRLUC* rhythm in the transfected cells is supported by surrounding tissues with normal circadian rhythms. In order to disrupt the intercellular signal transmission to the transfected cells, we exposed the frond to hypertonic medium to cause plasmolysis. We performed dual-color monitoring experiment on the fronds being subjected to NF medium with 500 mM mannitol (Figure 5). Plasmolysis of cells that were transfected with *CaMV35S::GFP-h* by particle bombardment was confirmed in the frond that was subjected to the hypertonic medium (Figure S5). The *AtCCA1::LUC*+ rhythm was observed in all analyzed cells in hypertonic conditions, although a period lengthening appeared (Figure 5A, 5C and Table S1). Interestingly though, the *CaMV35S::PtRLUC* rhythm disappeared in the same cells (Figure 5B, 5C and Table S1). Similar results were obtained for the circadian rhythmicity of each reporter in the single reporter experiments involving hypertonic treatment (Tables S2). The disappearance of the *CaMV35S::PtRLUC* rhythm in cells showing stable *AtCCA1::LUC*+ rhythm strongly suggests that the *CaMV35S::PtRLUC* rhythm is non-cell-autonomously generated with the support of surrounding tissues.

**Figure 5.**
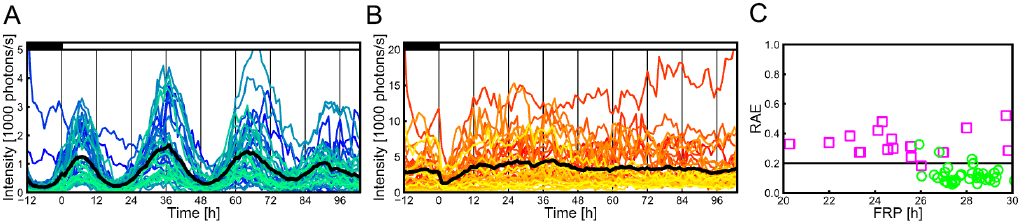
Cellular circadian properties of *AtCCA1::LUC*+ and *CaMV35S::PtRLUC* bioluminescence traces in a plant in hypertonic medium. A total of 30 cells in a frond on a hypertonic medium were analyzed for the dual-color reporters. (A and B) Cellular *AtCCA1::LUC*+ (A) and *CaMV35S::PtRLUC* (B) bioluminescence traces. A black line represents mean luminescence intensities. Close and open boxes represent light and dark conditions, respectively. (C) Free-running periods (FRPs) and the relative amplitude errors (RAEs) of individual cellular rhythms. Green circles and magenta squares represent *AtCCA1::LUC*+ and *CaMV35S::PtRLUC* bioluminescence traces, respectively. Cellular bioluminescence traces with RAE < 0.2 are defined as rhythmic.

## Discussion

In this study, we have demonstrated that the *CaMV35S::PtRLUC* bioluminescence rhythm is not governed by the clock gene circuit in the expressing cells but requires the support of surrounding tissues in the plant. The maintenance of high synchrony in the *CaMV35S::PtRLUC* rhythm between cells in the frond stands in contrast to the dyssynchronous nature of the *AtCCA1::LUC*+ rhythm, that is linked to the clock gene circuit (Figure 2F) (Muranaka and Oyama, 2016). This indicates distinct characteristics of cell-autonomous and non-cell-autonomous rhythms. Figure 6 shows a conceptual framework of generation of these rhythms in individual cells in the plant. In each cell, TTFL drives the cell-autonomous rhythms such as the *AtCCA1::LUC*+ rhythm. The non-cell-autonomous rhythms such as the *CaMV35S::PtRLUC* rhythm are driven by symplast/apoplast mediation. Circadian rhythms that are driven by changes in concentrations of ions and small molecules and the water flux potentially belong to this category. Circadian-regulated solute transporters of individual cells appear to generate these circadian rhythms (Haydon et al., 2011), while changes in the concentrations of these diffusive elements inevitably involve extracellular factors that are shared by the plant cells. These non-cell-autonomous rhythms can be driven at an organismal level. The rhythmicity at the organismal level depends on the integration of temporal information between cellular clocks. Therefore, a certain degree of synchrony between them is required to generate the non-cell-autonomous rhythms. This framework effectively explains the distinct characteristics of the *AtCCA1::LUC*+ and *CaMV35S::PtRLUC* rhythms. Diffusive elements such as luciferin, magnesium ions, protons, ATP, and oxygen that influence the luciferase enzyme activity (Van Leeuwen et al., 2000) may oscillate in a circadian manner in duckweed. In fact, a circadian rhythm of the absorption of ions such as potassium and magnesium ions, has been reported in the duckweed species *Lemna gibba* (Kondo, 1983; Kondo and Tsudzuki, 1978). It was also reported that *L. gibba* showed circadian rhythms in respiratory metabolism including the pentose phosphate pathway under constant light (Miyata and Yamamoto, 1969). The circadian rhythm at an organismal level should show higher synchrony between cells than the cell-autonomous rhythm. Thus, the *CaMV35S::PtRLUC* rhythm can be more synchronous than the *AtCCA1::LUC*+ rhythm in a frond. Notably the *CaMV35S::PtRLUC* rhythm is lost or is rendered unstable in a frond with cellular clocks in an asynchronous state (Watanabe et al., 2021).

**Figure 6.**
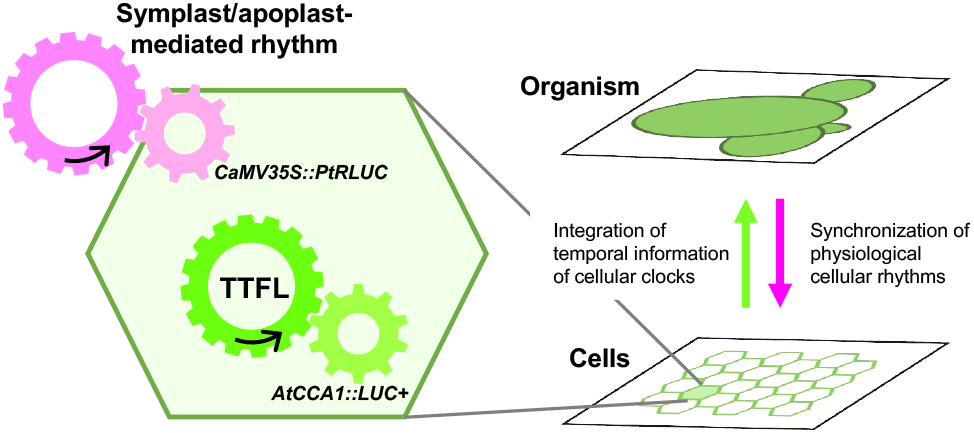
Conceptual framework of the generation of *AtCCA1::LUC*+ and *CaMV35S::PtRLUC* bioluminescence rhythms. The *AtCCA1::LUC*+ bioluminescence rhythm is generated by the TTFLs in individual cells. The *CaMV35S::PtRLUC* bioluminescence rhythm is a symplast/apoplast-mediated rhythm. Temporal information of cellular clocks is integrated to generate physiological symplast/apoplast-mediated rhythms at an organismal level. These physiological rhythms are synchronous between cells.

An important finding of this study is that physiological rhythms in individual cells are not always governed by their own cellular circadian clocks. These rhythms can stay stable in the plant even when the synchrony between the cellular clocks of its cells becomes low. This allows imprecision of individual cellular clocks and uneven circadian properties between cells. Indeed, in a *L. gibba* frond under constant light, each cellular clock showed an unstable FRP (SD for cycle-to-cycle variation: 2.45 h) and a relatively large variation of FRP was reported between cells (SD 1.11 h) (Muranaka and Oyama, 2016). The highly synchronous rhythm independent of the quality of individual cellular clocks would effectively work without the need to maintain the accuracy of each clock. Our results clearly indicate that the *CaMV35S::PtRLUC* rhythm was maintained in the cells in which the *AtCCA1::LUC*+ rhythm was disrupted by *ZTL* overexpression. This strongly suggests that the clock gene circuit is dispensable for the *CaMV35S::PtRLUC* rhythm in the cell. This does not necessarily mean that the gene circuit of the plant cells is completely dispensable for the *CaMV35S::PtRLUC* rhythm. The *ZTL* overexpression after transfection using particle bombardment was expected to disrupt the gene circuit only in the transfected cells, and most cells in the plant were not affected. These unaffected cells were probably responsible for the non-cell-autonomous rhythm at an organismal level. The stable transformation method has been established for *L. minor* (Chhabra et al., 2011; Ueno et al., 2022), and transgenic plants overexpressing *ZTL* would be an interesting model for studying the necessity of the clock gene circuit for the *CaMV35S::PtRLUC* rhythm. In a previous study, the circadian rhythms of bioluminescence reporters in roots (rootstocks) of *Arabidopsis* clock mutants were partially restored by grafting wild type shoot apex (Takahashi et al., 2015). In that case, it appears that the cellular circadian clock in the roots was restored. Therefore, the mechanism for generating the rhythm in *Arabidopsis* may differ from that of the *CaMV35S::PtRLUC* rhythm in duckweed. In fact, multiple mechanisms may exist to generate non-cell-autonomous circadian rhythms in plants.

Intercellular communication is an interesting aspect of the plant circadian system (McClung, 2019; Sorkin and Nusinow, 2021). Synchronization between neighboring cellular clocks and long-distance clock communication have been previously reported (Endo et al., 2014; Fukuda et al., 2012, 2007; Gould et al., 2018; Muranaka and Oyama, 2016; Takahashi et al., 2015; Ueno et al., 2022; Wenden et al., 2012). A non-cell-autonomous physiological rhythm with a high spatial synchrony can effectively function in the synchronization process between cellular clocks if the clock gene circuit is affected by the physiological rhythm in individual cells. Calcium is a potential mediator of this intercellular communication because cytosolic free calcium in plant cells is a physiological circadian output and is also involved in the regulation of the circadian clock (Johnson et al., 1995; Martí Ruiz et al., 2018). Interestingly, it has been suggested that the calcium rhythm and the *Cab* (*Lhcb*) gene expression rhythm are separately controlled for the determination of their FRPs and the oscillation generation by the clock gene circuit (Sai and Johnson, 1999; Xu et al., 2007). It is therefore worth investigating whether the calcium rhythm is non-cell-autonomous.

The mechanisms that generate the *CaMV35S::PtRLUC* bioluminescence rhythm are currently unknown (Watanabe et al., 2021). In the conceptual framework (Figure 6), physiological conditions associated with the bioluminescence reaction in the cells seem to post-transcriptionally mediate the rhythm. Circadian rhythms in the levels of ions and/or metabolites at an organismal level may bring about the bioluminescence rhythm (Haydon et al., 2013, 2011; Sorkin and Nusinow, 2021). These rhythms may not be the simple sum of the individual cellular rhythms in the plant because the FRP of the *CaMV35S::PtRLUC* rhythm is different from that of the *AtCCA1::LUC*+ rhythm (Table S1). Further studies of such non-cell-autonomous rhythms should improve the understanding the circadian systems in plants, which involve various cells with cell-autonomous circadian clocks.

## Materials and Methods

### Plant material and growth conditions

*Lemna minor* 5512 was maintained in NF medium with 1% sucrose under constant light conditions in a temperature-controlled room (25 ± 1 °C), as previously described.(Muranaka et al., 2015) The white light (~20 μE m^-2^s^-1^) was generated by fluorescent lamps (FLR40SEX-W/M/36-HG, NE, Japan). Plants were grown on 60 ml of the medium in 200-ml Erlenmeyer flasks plugged with cotton. New stock cultures were made weekly, and well-grown plants were used for the experiments. To entrain the circadian rhythm, plants were cultured under at least three 12 h light/12 h dark cycles in an incubator (MIR-153, Panasonic, Japan) before starting the experiments.

### Luciferase reporters

As circadian bioluminescent reporters, we used *pUC-AtCCA1::intron-LUC*+ (*AtCCA1::LUC*+), and *pUC18-CaMV35S::PtRLUC* (*CaMV35S::PtRLUC*), which have been described previously(Watanabe et al., 2021). PtRLUC is a red-shift mutant (S246H/H347A) of Emerald luciferase (ELuc) from Brazilian click beetle (*Pyrearinus termitilluminans*).(Nishiguchi et al., 2015)

### Clock-gene overexpression effectors

*pUC18-CaMV35S::LmZTL, pUC18-CaMV35S::LmLUX*, and *pUC18-CaMV35S::LmELF3* were constructed as follows. Coding regions of *LmZTL, LmLUX*, and *LmELF3* from *Lemna minor* 5512 were amplified using PCR with the following primer sets: CgcggccgcCACCATGGAGTGGAATAGTGATTCG and CggcgcgccTTAAATGAGCTCCGAGAAGCC for *LmZTL*, CgcggccgcATGGCGGACGGTGATGGCGG and CggcgcgccTTATTTGTAACCGGGAGAAACATGG for *LmLUX*, CgcggccgcATGAAGAGAGGGAGAGAGGAAGATAAG and CggcgcgccCTACAGCGGATCGTATCCCTG for *LmELF3*. Primer sequences for each gene were designed based on the genome sequence of *Lemna minor* 5500.(Van Hoeck et al., 2015) The small-letter 8-b sequence in each primer sequence represents a restriction enzyme recognition site (*Not1* for forward primer and *Asc*1 for reverse primer). cDNA that was synthesized from total RNA of *L. minor* 5512 using ReverTra Ace (TOYOBO, Japan) was used as the template for the PCR. The total RNA was extracted from the whole plant using NucleoSpin RNA (Takara, Japan). Sequence data of *LmZTL_1* (LC708291), *LmLUX_1* (LC708289) and *LmELF3_1* (LC708287) are posted at DDBJ. The amino acid sequences of LmZTL and LmELF3 are identical in the strains 5500 and 5512. Compared with the strain 5500, the amino acid sequence of LmLUX in strain 5512 contains four amino acid substitutions and an insertion of two amino acids. The amplicon of each gene was cloned into *pENTR-D-TOPO* (Invitrogen, MA, USA) with an additional multi-cloning site. Using the LR reaction, the coding region in this plasmid was integrated into *pUC18-CaMV35S-attR1-attR2-NosT*, in which the *att*R1-*att*R2 sequence (Invitrogen) was located between the *CaMV35S* promoter and the Nos terminator.

### Particle bombardment and bioluminescence monitoring

Reporter constructs were introduced into *L. minor* plants by particle bombardment as described previously.(Muranaka et al., 2015) In both single- and dual-reporter experiments, the amounts of plasmid DNA coding for *AtCCA1::LUC*+ and *CaMV35S::PtRLUC*, were 2 μg, and 0.2 μg, respectively. In co-transfection experiments, 1 μg overexpression effector of a clock gene was added to the reporter(s).

Dual-color bioluminescence monitoring at a single cell level was carried out as described previously.(Watanabe et al., 2021) In short, the imaging system consisted of an EM-CCD camera (ImagEM C9100-13; Hamamatsu Photonics, Japan) with a lens (VS-50085/C 0.85/50mm; VS Technology, Japan), a handmade filter changer with a rotation motor (ARS-4036-GM; Chuo Precision Industrial Co., Japan), a handmade motor-driven stage with rotation and z-axis stages (ARS-6036-GM and ALV-902-HP; Chuo Precision Industrial Co., Japan), and an illuminating system with an end-emitting fiber optics guiding LED white light (approximately 30 μE m^-2^ s^-1^; RFB2-20SW; CCS Inc, Japan). These devises were set in a lightproof box in an incubator (25 ± 1°C). A green-pass filter (PB0530/080; ⌀49.7 mm; Asahi Spectra, Japan) and red-pass filter (PB0630/040; ⌀49.7 mm; Asahi Spectra) were placed in slots in the filter changer, and one slot was left empty to capture unfiltered images. By controlling this imaging system using PC software (HOKAWO; Hamamatsu Photonics), a series of bioluminescence images for a sample dish were automatically captured every 1 h. Based on the intensities of the filtered bioluminescence of a luminous cell expressing both the reporters, the bioluminescence intensity of each reporter at every time point was calculated. The same imaging protocol was used for single-reporter experiments. Bioluminescence monitoring in the hypertonic conditions (NF medium with 1% sucrose and 500 mM mannitol) was carried out using a plant that was transferred from the normal medium conditions just before the start of the monitoring.

### Fluorescence imaging

A fluorescent reporter (*CaMV35S::GFP-h*)(Nakano et al., 2009) was transfected into the duckweed plants, and GFP fluorescence-positive cells were observed under a confocal microscope (LSM510-META; Carl Zeiss, https://www.zeiss.com) as described previously.(Muranaka et al., 2013)

### Time series analysis

The criteria for luminous spots that were used for time series analysis were the same as those previously described, except for a change in the lower limit of the ratio of luminescence intensities of a 24 h moving average between *AtCCA1::LUC*+ and *CaMV35S::PtRLUC* (from 1/20 to 1/100).(Watanabe et al., 2021) The FRPs and RAEs of the bioluminescence traces were estimated using the fast Fourier transform nonlinear least squares method, as described previously.(Watanabe et al., 2021) The peak times of bioluminescence traces were estimated by local quadratic curve fitting, as described previously.(Watanabe et al., 2021)

Phase *θ*(*t*) of the bioluminescence rhythms *x*(*t*) was defined as follows

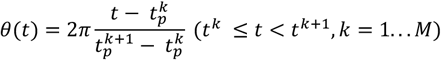

where 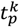 represents the occurrence time of the *k*^th^ peak of *x*(*t*) and *M* represents the total number of peaks. Data analysis codes are available at https://github.com/ukikusa/circadian_analysis. Timeseries data of cellular bioluminescence will be deposited at KURENAI (https://repository.kulib.kyoto-u.ac.jp/dspace/).

### Synchronization index

To assess the synchrony between cellular rhythms within a plant, the SI, which is known as the order parameter in the Kuramoto model, was calculated at each time point as follows

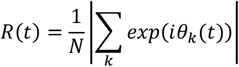

where *θ_k_*(*t*) is the phase of the *k*^th^ cell at time *t*, and *N* is the number of cells.

The SI at a given time was calculated if the ratio of cells of which the phase was defined was more than 80% of rhythmic cells. For dual reporter experiments, cells showing both rhythms were analyzed.

## Acknowledgments

We thank Dr. Hiroshi Kori for fruitful discussions and useful advice. This research was supported by the Japan Society for the Promotion of Science KAKENHI [grant numbers JP21J23250 (E.W.), JP19J23441(S.N.), JP16H06864 (T.M.), JP20K06342 (S.I.), JP17KT0022 (T.O.) and JP19H03245(T.O.)], Japan Science and Technology Agency (JST), JST ALCA (JPMJAL1108, T.O.), and JST SATREPS (JPMJSA2004, T.M., S.I., T.O.). (Watanabe et al., 2021)

---

**Figure S1.**
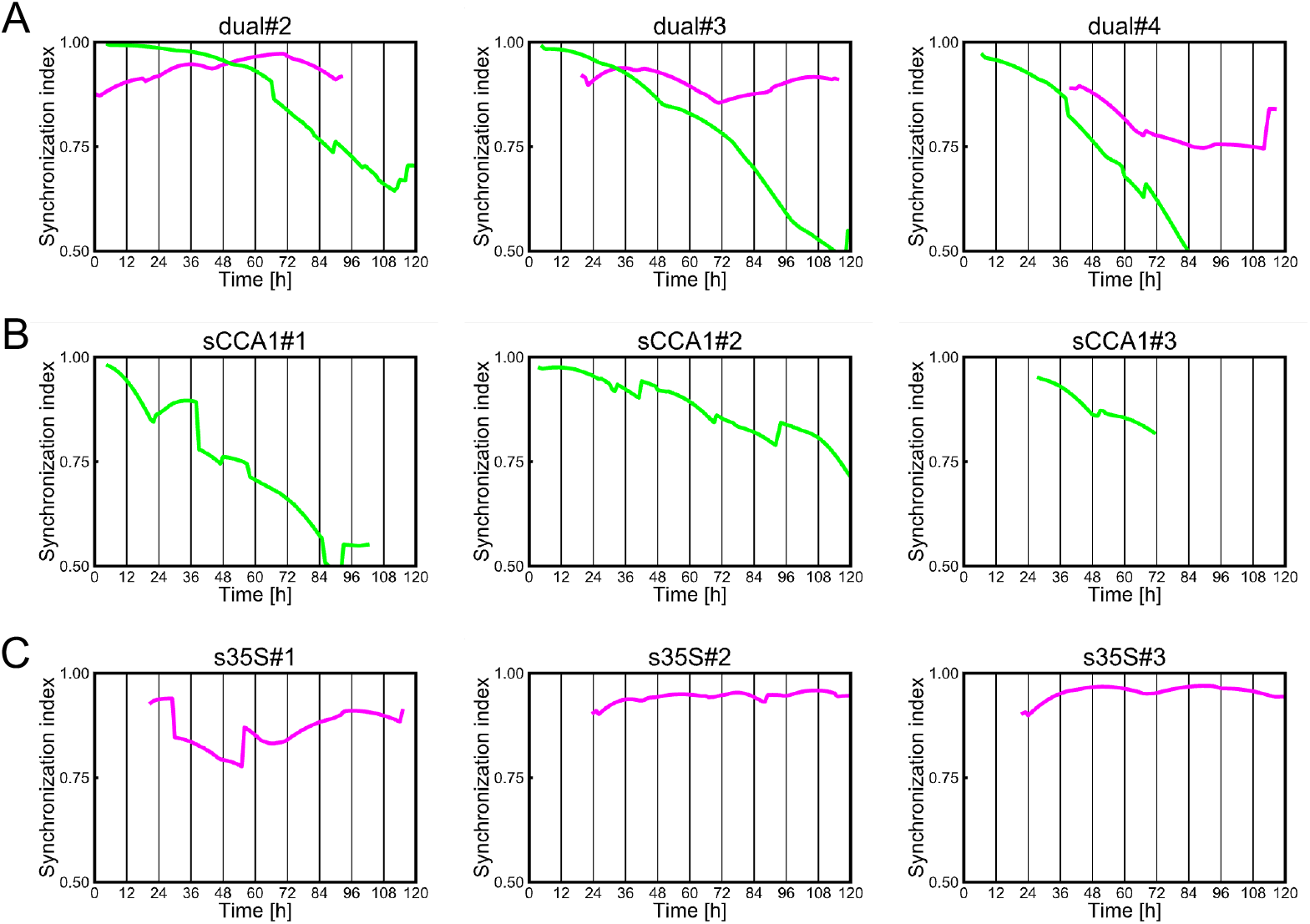
Temporal changes in the synchronization index (SI) of *AtCCA1::LUC*+ (green) and *CaMV35S::PtRLUC* (magenta) rhythms in dual and single reporter experiments. (A) Dual-reporter experiments (dual#2-#4), (B) Single-reporter experiments for the *AtCCA1::LUC*+ rhythm (sCCA1#1-#3), (C) Single-reporter experiments for the *CaMV35S::PtRLUC* rhythm (s35S#1-#3).

**Figure S2.**
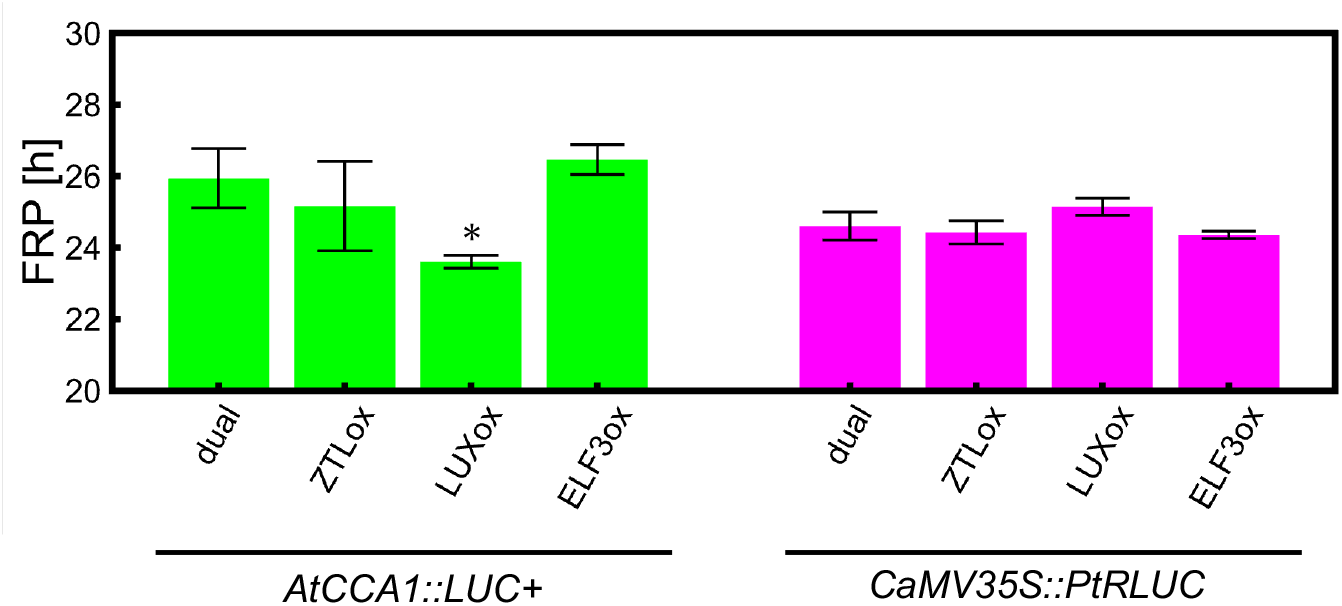
Free running periods (FRPs) of *AtCCA1::LUC*+ and *CaMV35S::PtRLUC* bioluminescence rhythms in overexpression experiments. FRPs of cellular bioluminescence rhythms in the control (dual; dual#1-#4), *LmZTL* (ZTLox; dual-ZTLox#1-#3), *LmLUX* (LUXox; dual-LUXox#1-#3) and *LmELF3* (ELF3ox; dual-ELF3ox#1-#3) overexpression experiments. Data are expressed as mean ± SE. Asterisk denotes statistically significant difference (Welch’s *t* test, *p* < 0.05).

**Figure S3.**
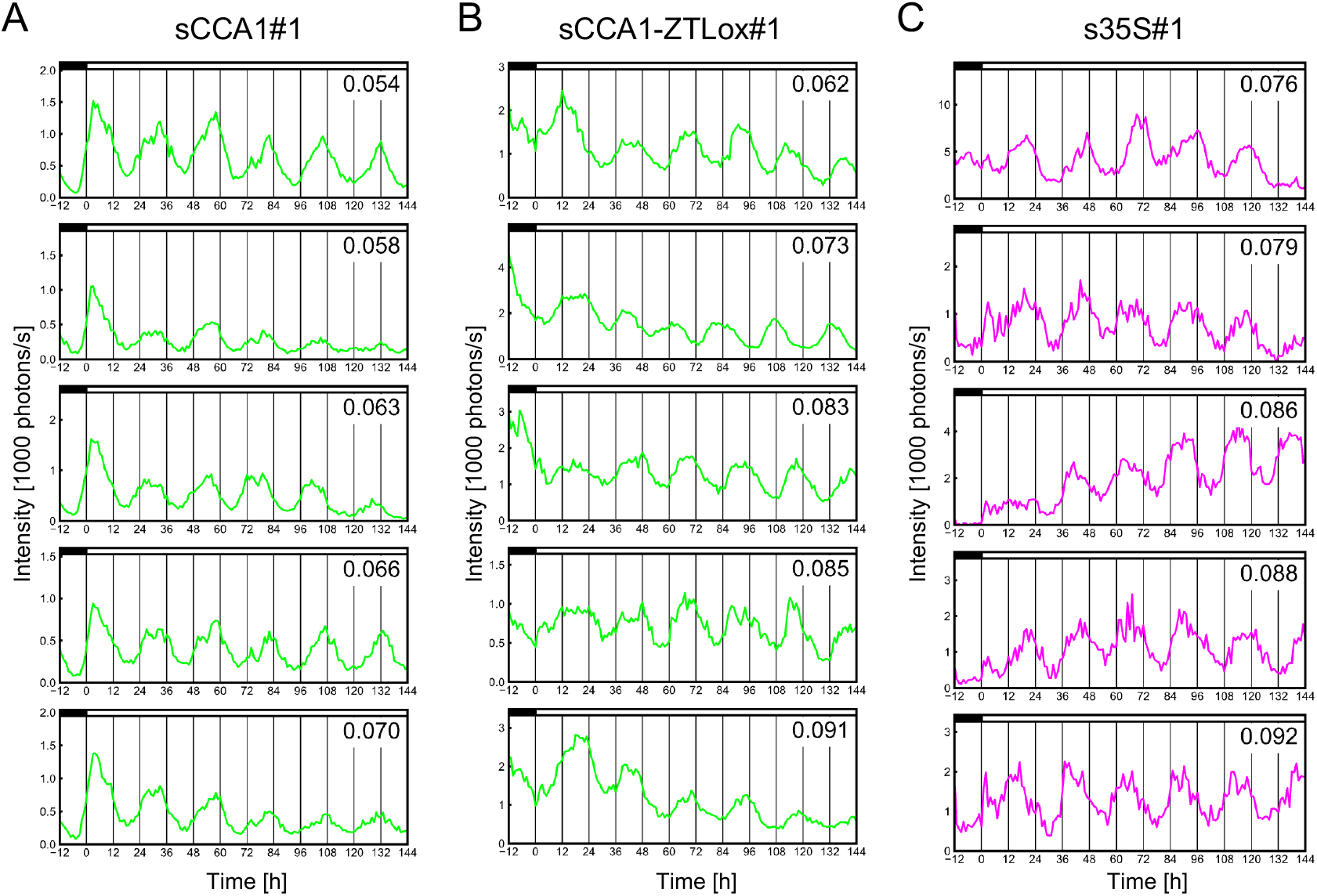
Cellular *AtCCA1::LUC*+ bioluminescence rhythms in an *LmZTL* overexpression experiment with single-reporter. Examples of bioluminescence rhythms of individual cells in single-reporter experiments: sCCA1#1 (A), sCCA1-ZTLox#1 (B), and s35S#1 (C). The cellular bioluminescence rhythms with the five lowest relative amplitude error (RAE) values in each experiment are shown in ascending order. The RAE is indicated in each graph. Close and open boxes represent light and dark conditions, respectively.

**Figure S4.**
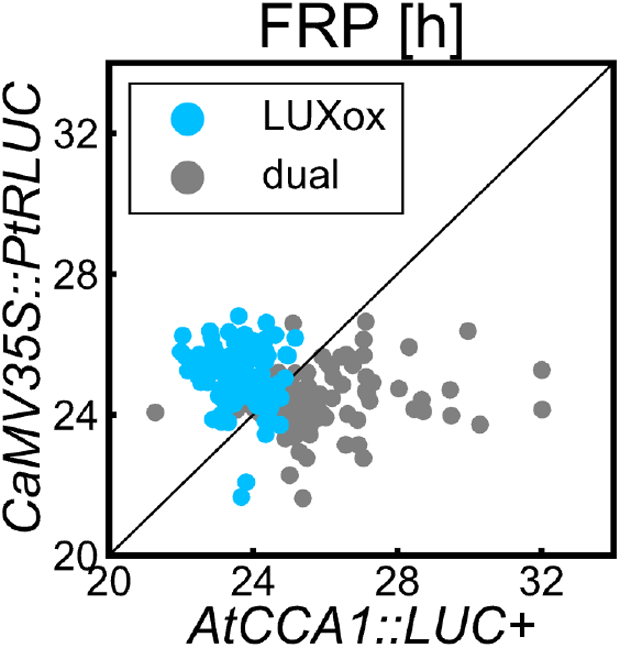
A scatterplot of the free-running periods (FRPs) between *AtCCA1::LUC*+ and *CaMV35S::PtRLUC* rhythms in dual-reporter experiments with or without *CaMV35S::LmLUX*. FRPs of the 132 cells showing the circadian rhythms of both reporters in the three overexpression experiments (dual-LUXox#1-#3) are plotted with blue dots. FRPs of the 116 cells showing the circadian rhythms of both reporters in the four experiments without the effector (dual#1-#4) are plotted with gray dots (the same plots as those in Figure 3B). The dashed line represents where the FRPs are identical in both rhythms.

**Figure S5.**
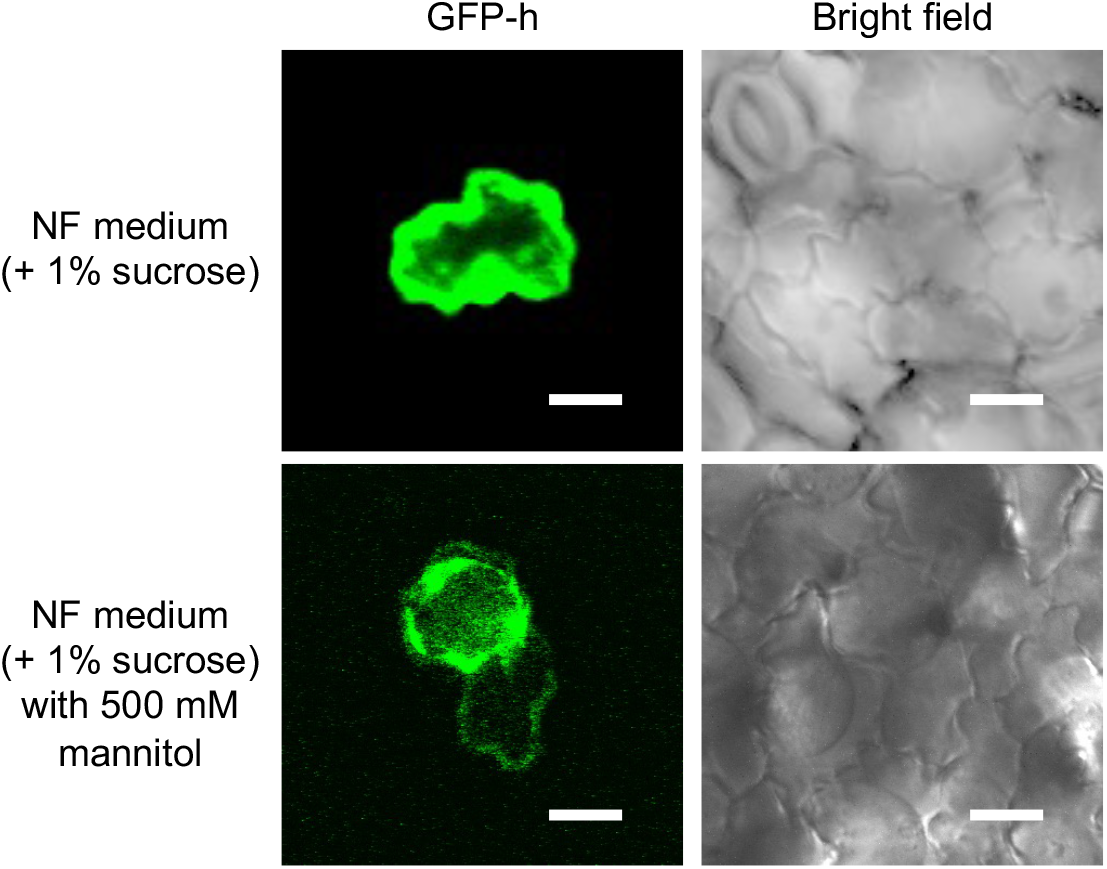
Plasmolysis of cells under hypertonic conditions. GFP fluorescence images of cells that were transfected with an endoplasmic reticulum (ER) localized GFP reporter (*CaMV35S::GFP-h*) and bright field images are shown. An epidermal cell of an *L. minor* plant in NF medium with 1% sucrose and that in NF medium with 1% sucrose and 500 mM mannitol are shown. Bars represent 10 μm.

**Table S1.**
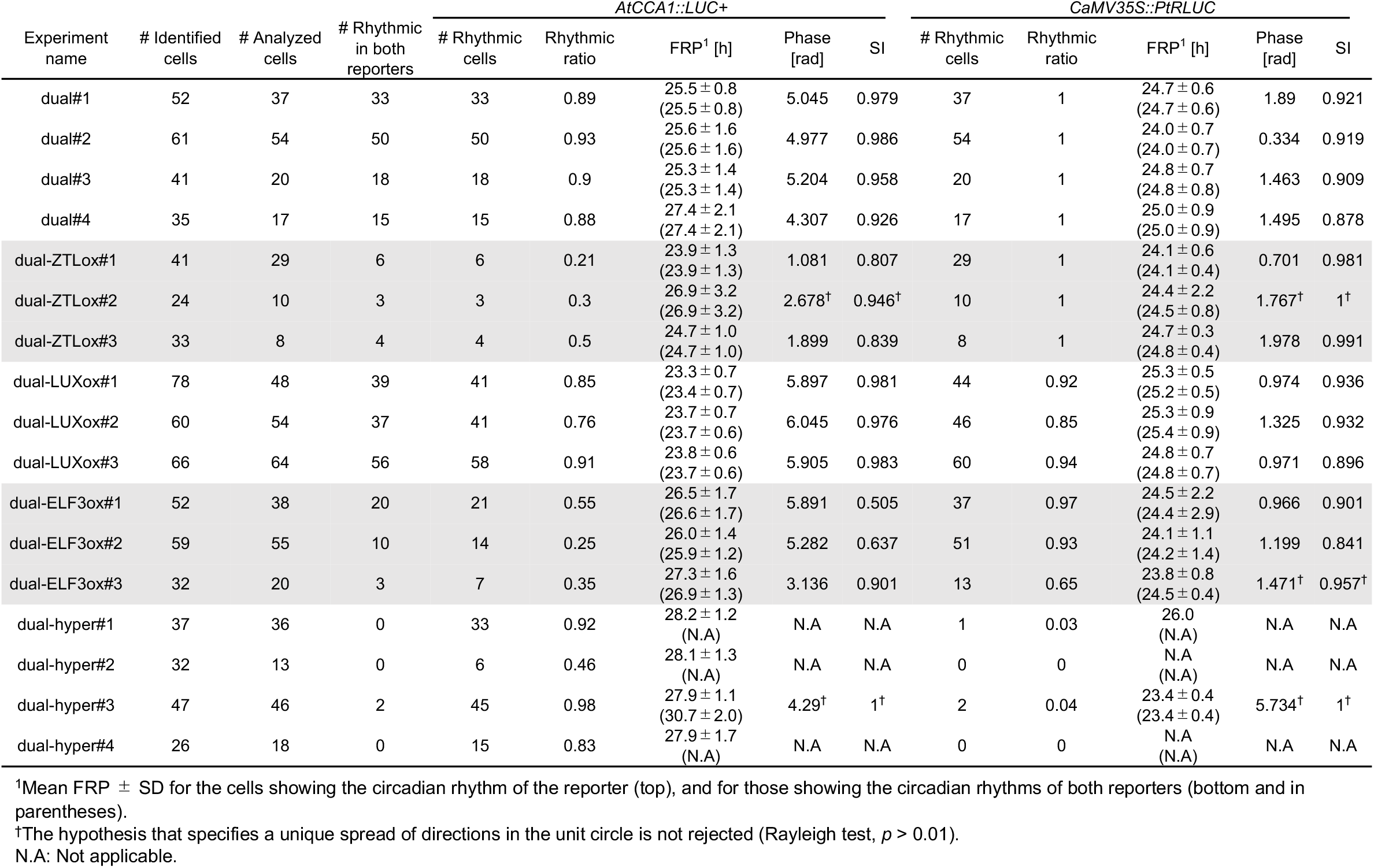
Summary of the quantitative analysis of cellular bioluminescence rhythms in dual reporter experiments

**Table S2.** Summary of the quantitative analysis of cellular bioluminescence rhythms in single reporter experiments

| Experiment name | # Identified cells | # Analyzed cells | *AtCCA1::LUC+* |  |  |  |  | *CaMV35S::PtRLUC* |  |  |  |  |
| --- | --- | --- | --- | --- | --- | --- | --- | --- | --- | --- | --- | --- |
|  |  |  | # Rhythmic cells | Rhythmic ratio | FRP<sup>1</sup> [h] | Phase [rad] | SI | # Rhythmic cells | Rhythmic ratio | FRP<sup>1</sup> [h] | Phase [rad] | SI |
| sCCA1#1 | 21 | 21 | 19 | 0.9 | 25.4 ± 1.3 | 5.558 | 0.865 | - | - | - | - | - |
| sCCA1#2 | 43 | 42 | 42 | 1 | 24.6 ± 0.8 | 5.654 | 0.954 | - | - | - | - | - |
| sCCA1#3 | 18 | 15 | 10 | 0.67 | 25.1 ± 1.0 | 5.284 | 0.983 | - | - | - | - | - |
| s35S#1 | 29 | 22 | - | - | - | - | - | 19 | 0.86 | 23.4 ± 2.7 | 1.087 | 0.937 |
| s35S#2 | 47 | 47 | - | - | - | - | - | 45 | 0.96 | 24.6 ± 0.5 | 0.706 | 0.905 |
| s35S#3 | 53 | 52 | - | - | - | - | - | 50 | 0.96 | 24.7 ± 0.6 | 0.819 | 0.899 |
| sCCA1-ZTLox#1 | 35 | 35 | 17 | 0.49 | 23.7 ± 2.4 | 1.847 | 0.819 | - | - | - | - | - |
| sCCA1-ZTLox#2 | 33 | 33 | 22 | 0.67 | 24.5 ± 0.6 | 1.774 | 0.906 | - | - | - | - | - |
| s35S-ZTLox#1 | 37 | 37 | - | - | - | - | - | 35 | 0.95 | 24.1 ± 0.7 | 0.384 | 0.886 |
| s35S-ZTLox#2 | 38 | 38 | - | - | - | - | - | 36 | 0.95 | 24.7 ± 0.9 | 0.297 | 0.953 |
| sCCA1-LUXox#1 | 34 | 32 | 31 | 0.97 | 23.6 ± 0.8 | 5.975 | 0.932 | - | - | - | - | - |
| sCCA1-LUXox#2 | 26 | 23 | 21 | 0.91 | 23.6 ± 1.2 | 5.713 | 0.972 | - | - | - | - | - |
| s35S-LUXox#1 | 51 | 51 | - | - | - | - | - | 51 | 1 | 24.6 ± 0.5 | 1.118 | 0.923 |
| s35S-LUXox#2 | 44 | 44 | - | - | - | - | - | 41 | 0.93 | 25.0 ± 0.9 | 1.342 | 0.905 |
| sCCA1-ELF3ox#1 | 37 | 37 | 25 | 0.68 | 27.0 ± 3.4 | 5.147 | 0.53 | - | - | - | - | - |
| s35S-ELF3ox#1 | 57 | 56 | - | - | - | - | - | 55 | 0.98 | 22.8 ± 1.0 | 0.881 | 0.856 |
| s35S-ELF3ox#2 | 68 | 68 | - | - | - | - | - | 53 | 0.78 | 22.3 ± 1.4 | 1.269 | 0.872 |

<sup>1</sup>Mean FRP ± SD for rhythmic cells.

